# Epigenetic landscape reorganization and reactivation of embryonic development genes are associated with malignancy in IDH-mutant astrocytoma

**DOI:** 10.1101/2024.03.19.585212

**Authors:** Santoesha A. Ghisai, Levi van Hijfte, Wies R. Vallentgoed, C. Mircea S. Tesileanu, Iris de Heer, Johan M. Kros, Marc Sanson, Thierry Gorlia, Wolfgang Wick, Michael A. Vogelbaum, Alba A. Brandes, Enrico Franceschi, Paul M. Clement, Anna K. Nowak, Vassilis Golfinopoulos, Martin J. van den Bent, Pim J. French, Youri Hoogstrate

**Author notes:** These authors contributed equally to this work (Levi van Hijfte, Wies R. Vallentgoed, Martin J. van den Bent, Pim J. French, Youri Hoogstrate).

## Abstract

**Purpose:** Accurate grading of IDH-mutant gliomas defines patient prognosis and guides the treatment path. Histological grading is however difficult and, apart from *CDKN2A/B* homozygous deletions in IDH-mutant astrocytomas, there are no other objective molecular markers used for grading.

**Experimental Design:** RNA-sequencing was conducted on primary IDH-mutant astrocytomas (n=138) included in the prospective CATNON trial, which was performed to assess the prognostic effect of adjuvant and concurrent temozolomide. We integrated the RNA-sequencing data with matched DNA-methylation and NGS data. We also used multi-omics data from IDH-mutant astrocytomas included in the TCGA dataset and validated results on matched primary and recurrent samples from the GLASS-NL study.

**Results:** We used the DNA-methylation profiles to generate a Continuous Grading Coefficient (CGC) that is based on classification scores derived from a CNS-tumor classifier. We found that the CGC was an independent predictor of survival outperforming current WHO-CNS5 and methylation-based classification. Our RNA-sequencing analysis revealed four distinct transcription clusters that were associated with i) an upregulation of cell cycling genes; ii) a downregulation of glial differentiation genes; iii) an upregulation of embryonic development genes (e.g. *HOX*, *PAX* and *TBX*) and iv) an upregulation of extracellular matrix genes. The upregulation of embryonic development genes was associated with a specific increase of CpG island methylation near these genes.

**Conclusions:** Higher grade IDH-mutant astrocytomas have DNA-methylation signatures that, on the RNA level are associated with increased cell cycling, tumor cell de-differentiation and extracellular matrix remodeling. These combined molecular signatures can serve as an objective marker for grading of IDH-mutant astrocytomas.

## Introduction

The fifth edition of the World Health Organization (WHO) classification of central nervous system tumors (WHO CNS5) incorporates molecular markers to classify distinct entities (1). Grading of CNS tumors is nevertheless mainly reliant on histopathological criteria. Marking these features in IDH-mutant astrocytomas is difficult, subjective and therefore prone to inter-observer variability (2). Apart from *CDKN2A/B* homozygous deletion (HD), there are no molecular markers that currently aid grading of IDH-mutant astrocytomas. However, precise tumor grading is crucial for improved patient surveillance and treatment. The latter was emphasized by results from the recently published INDIGO trial showing strong clinical efficacy of the IDH1/2 inhibitor vorasidenib in grade 2 IDH-mutant glioma (3).

Recent advances in molecular profiling resulted in the development of DNA methylation-based classification, which offers an objective alternative approach to histology-based grading (4,5). DNA-methylation-based classification has been integrated into WHO CNS5 as diagnostic criteria for several CNS tumor types (6). The “CNS tumor classifier” uses a reference cohort of over 2,800 CNS tumors and predicts tumor (sub)classes that are determined on unsupervised methylation-intrinsic clusters (5). Apart from classification, the CNS tumor classifier stratifies IDH-mutant astrocytomas into two subclasses, low-(A_IDH_LG) and high-grade (A_IDH_HG), with distinct prognoses (7).

In earlier work, we observed that the variation in DNAm profiles appears to be continuous and associated with tumor malignancy (7). Here we used genome-wide DNA-methylation (DNAm) data from samples included in the multicenter CATNON randomized phase III clinical trial on anaplastic gliomas with absence of combined loss of the 1p and 19q chromosomal arms (1p/19q codeletion) (8) to construct a Continuous Grading Coefficient (CGC). We identified that increased malignancy is associated with glial dedifferentiation, upregulation of extracellular matrix (ECM) genes and increased expression of cell cycling genes. Also, while acknowledging continuity of tumor grading, our cut-off points for the CGC showed a more precise method of classification compared to WHO CNS5 and results from the ‘CNS tumor classifier’.

## Materials and Methods

### Study cohort

We included subjects from the CATNON (8), TCGA (9) and GLASS-NL (10) studies. In the non-blinded randomized CATNON trial patients aged older than 18 years with newly diagnosed 1p/19q non-codeleted anaplastic gliomas and a WHO performance score of 0-2 were included (8). This multicenter study included patients from 137 institutes across Australia, Europe and North America. The 2×2 factorial design compared radiotherapy alone or radiotherapy combined with adjuvant temozolomide to those receiving radiotherapy and concurrent temozolomide, or radiotherapy with both concurrent and adjuvant temozolomide (December 2007–September 2015). The primary endpoint was overall survival (OS), measured from the day of randomization until death or last follow-up. More detailed information regarding endpoints, inclusion and exclusion criteria are described elsewhere (8). Central histology review of samples from the CATNON study detected necrosis and/or microvascular proliferation in some cases, leading to their re-classification as WHO grade IV (WHO CNS4) (11). In the present study, we specifically analyzed the subset of IDH-mutant samples from the CATNON trial (median follow-up time: 54.5 months, Supplementary Methods). For this post-hoc study, sample size was restricted by the original trial design. The GLASS-NL study systematically investigated longitudinal changes in patients with an initial diagnosis of IDH-mutant astrocytoma (grade II/III, WHO CNS4) who had undergone at least two resections with more than 6 months intervals in between (10). This multicenter (n=3) study included patients treated in The Netherlands. Histopathological diagnoses were reassessed by a dedicated neuropathologist. Treatment regimens in this cohort were heterogeneous and involved either radiotherapy or chemotherapy alone, or a combination of both, where progression was not consistently followed by surgical resection. Treatment regimen in the TCGA cohort were not controlled by the study design and were based on standard clinical practices and personalized patient assessments. To explore mechanisms associated with increased malignancy prior to post-surgery treatment, we exclusively utilized molecular data from initial resections for the CATNON and TCGA datasets. The GLASS-NL dataset was partitioned into primary and recurrent subsets.

### RNA-sequencing

For this study, we RNA-sequenced material of tumor resections of the randomized phase 3 CATNON trial (NCT00626990). Written informed consent was obtained from all participants in accordance with institutional and national ethical guidelines. Tumor tissue samples from IDH-mutant astrocytomas in the CATNON dataset (n=306) were obtained from adult patients through the European Organisation for Research and Treatment of Cancer (EORTC) study 26053-22054. RNA was extracted from formalin-fixed paraffin-embedded (FFPE) tissue blocks, stored at room temperature, using the RNeasy FFPE kit according to the manufacturer’s protocol (#73504, QIAGEN, Germany). Material was isolated by a technician not involved in the data analysis and only aware of pseudonymized storage identifiers. Library quality control included RNA integrity analysis using the Fragment Analyzer (Agilent Technologies) and RNA concentration quantification with the Qubit fluorometer (Invitrogen). Samples with compromised integrity and RNA concentrations below 0.5 nM were excluded (n=123) (GenomeScan BV, Leiden, The Netherlands). RNA-sequencing (n=183 samples) was performed on an Illumina NovaSeq 6000 with 150 bp paired-end reads including Unique Molecular Identifier (UMI) tags (GenomeScan BV, Leiden, The Netherlands). Sample exclusion criteria and processing of FASTQ files into alignments and read counts are described in the Supplementary Methods. In total, 148 samples passed stringent quality control from which we only included first resections with matching methylation data (n=138).

### Molecular and clinical data collection

For the CATNON dataset Copy Number Variation (CNV) data were extracted from earlier generated Infinium EPIC DNAm data (7), where HD and amplifications were defined according to previously described criteria (7). Detailed methodologies and protocols used for obtaining our previously generated Infinium EPIC DNAm data are described in the Supplementary Methods (7). *IDH1/2* mutation status was determined from 1 of 2 glioma-tailored next-generation sequencing (NGS) panels (7). The 1p/19q codeletion status had been determined by local testing or at central review. TCGA clinical, RNA-sequencing and Infinium 450k DNAm data were downloaded from the TCGA-LGG and TCGA-GBM projects (9) through the Genomic Data Commons (GDC) portal. CNV calls were obtained from cBioPortal (www.cbioportal.org). Annotations for the *IDH1/2* mutation and chromosome 1p/19q codeletion status were obtained from literature (9). For the GLASS-NL dataset, raw Infinium EPIC DNAm and RNA-sequencing data were collected from the public data repository with accession number XXX (upload to EGA pending). *IDH1/2,* CNV and 1p/19q codeletion status were obtained from earlier work (12).

### CNS tumor classifier

Methylation arrays were batch analyzed for all datasets using the Heidelberg CNS tumor classifier (“report_website_mnp_brain_v12.8_sample (Version 1.1)”, https://www.molecularneuropathology.org) (5). To facilitate throughput of the excessive size of the datasets, classifications were batch executed, and results were batch-downloaded using our *pymnp* API (https://github.com/yhoogstrate/pymnp). The CGC were calculated from the predictBrain_v12.8 *_scores_cal.csv files:

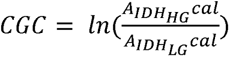

Hereby *cal* refers to the calibrated score, which is determined through a probability calibration model. This model transforms the raw scores into probabilities reflecting the confidence in class assignment (5). Methylation-based CNV estimations were obtained from the “cnvp_v5.2/*.bins.igv” files. We used a threshold of +/- 0.10 log2 intensity value for loss and gain, where samples exceeding the cut-off value of 350 Mb were categorized as having a high CNV load (13).

### CGC association with copy-number events

For each genomic bin from the “cnvp_v5.2/*.bins.igv” file, we acquired a measure of malignancy per individual copy number event by conducting a Wilcoxon signed-rank test with FDR-adjustment on the CGC. Associations of the CGC with copy number events were calculated for the amplification and HD status independently. We used log2 intensity differences of +0.35 and −0.415 as cut-off for amplification and homozygous deletion, as proposed in literature (13). These tests were performed independently for each dataset, ensuring dataset-specific associations. Results were presented as log10(p-values) for losses and -log10(p-values) for gains.

### DNA-methylation Analysis

Raw DNAm data of matched methylation and RNA samples were preprocessed as described earlier (Supplementary Methods) (7). For each probe, the methylation intensity values from the methylated and unmethylated channels were converted to obtain beta-values (beta = methylated intensity/(methylated+unmethylated intensity)) (14). These values were then transformed to M-values, which are logit-transformed beta-values, to improve homoscedasticity (15). Linear regression modelling on the M-values was performed for Differential Methylation Probe (DMP) analysis. We excluded probes targeting the sex chromosomes and those lacking a UCSC RefGene annotation (Illumina Human Methylation EPIC annotation 1.0 B5). DMP analysis was done using limma’s lmfit function for model fitting and probe-wise fold changes of the M-values were estimated using empirical Bayes moderation (limma::eBayes) of the standard errors (16). Probes with an absolute log2-transformed fold change (LFC) larger than 0.5 (FDR-adjusted p-value<0.01) were considered significantly differentially methylated.

### mRNA expression analysis

Lowly expressed genes with an average read count of less than three reads per sample were omitted. Also, only genes with *gencodev34* gene type annotation “protein coding” and “lncRNA” were included. DESeq2 (17) was used to estimate variance stabilized transformed (VST) normalized read counts for each dataset separately. Differential Gene Regression (DGR) analysis was performed using the Wald test (17) where the CGC was used as continuous condition. Genes with an absolute LFC higher than 0.5 and FDR-adjusted p-value smaller than 0.01 were considered significantly differentially expressed. Recursive-based correlation clustering on the differentially expressed genes was performed using the recursiveCorPlot package (18,19). Clusters with 50 genes or fewer were excluded.

### Correlation between RNA expression and DNAm

Probe-level M-values were transformed to gene-level M-values by computing the median M-values of all probes linked with a gene as defined by the UCSC RefGene annotation (Illumina Human Methylation EPIC annotation 1.0 B5). Genes that were also present in the processed RNA-sequencing data were then included. The overall correlation between the DMP (log2FoldChange M-value) and DGR (log2FoldChange normalized expression value) analyses were visualized by comparing the z-statistics per gene (log2FoldChange/standard error log2FoldChange).

### vanHijfte_2023 single-nucleus RNA-sequencing dataset and analysis

For single-nucleus RNA-sequencing (snRNA-seq) analysis of primary-recurrent astrocytoma samples, our in-house vanHijfte_2023 dataset was used. According to the WHO CNS4 classification of gliomas (11), primary samples were classified as low-grade and progressed to high-grade in the matched recurrences. Data preprocessing and normalization were performed as described previously (12). Briefly, nuclei doublets were filtered out using the *scDblFinder* package (20). For every sample, a lower limit of unique reads was defined to exclude low complex measurements. Additionally, nuclei with a mitochondrial read fraction of >0.1 were removed. Next, data was normalized using the SCTransform V2 function from the SCTransform package (21). All samples were integrated into one dataset using the reciprocal Principal Component Analysis (PCA) method as integrated in the Seurat package (v4) (22). Cell types were annotated using markers for normal cell types and malignant transcriptomic cell states (23–25). Enrichment of marker gene sets was assessed using the gene function described previously (23). In short, for a gene set G, a reference gene set R was created. First, all genes were separated into 30 equal-sized bins according to expression level. Next, for every gene in G, 100 random genes from its corresponding bin were added to R, resulting in R being 100x larger than G. The difference in average expression for G and R is calculated for every nucleus to provide an enrichment score.

### Statistical analysis and visualization

All statistical analyses were performed in R programming language (v.4.2.2). Survival analysis was performed using the *survival* package (v3.5-7). We examined OS as clinical endpoint for all survival analyses. Overall survival probabilities were calculated using the Kaplan-Meier estimator and log-rank tests were used to compare survival curves. Cox proportional hazard (PH) regression modelling was performed to determine hazard ratios (HR) for univariable and multivariable analyses, where significance was assessed using the likelihood ratio test. For multivariable analyses, comparisons were made against markers already included in WHO CNS5 (*CDKN2A/B* HD and microvascular proliferation/necrosis). Cut-off points for the CGC were determined unsupervised with respect to clinical end points by assessing the rate of change of 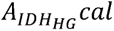 per unit increase in CGC. The CGC-based “medium-grade” group comprised 95% of the total cumulative rate. Cut-off values were first defined on CATNON and independently validated on TCGA and GLASS-NL. Gene set enrichment analysis (GSEA) was performed using the *clusterProfiler* package (v4.6.2). Molecular Signatures Database (MSigDB) human gene set collections (C2 and C5) were extracted using the *msigdbr* package (v.7.5.1) (26). Plots were generated with the *tidyverse* package (v2.0.0). Survival forest plots of CoxPH models were visualized using the *survminer* package (v0.4.9).

## Results

### Sample Cohort

To investigate biological mechanisms underlying malignancy of IDH-mutant astrocytomas, four large multi-domain and multi-center omics datasets were leveraged: CATNON, TCGA, GLASS-NL primary (GLASS-NL-P) and GLASS-NL recurrent (GLASS-NL-R) (8–10). We obtained DNAm (CATNON: n=430, TCGA: n=256, GLASS-NL-P: n=98, GLASS-NL-R: n=137) and DNA-sequencing (CATNON: n=424, TCGA: n=253, GLASS-NL-P: n=97, GLASS-NL-R: n=133) data for each of the studies. RNA-sequencing of primary IDH-mutant astrocytomas included in the CATNON trial was successfully performed for 138 samples and extended with transcriptomic data from the TCGA (n=247), GLASS-NL-P (n=65) and GLASS-NL-R (n=102) datasets.

The prevalence of grade IV tumors was notably higher in the CATNON dataset, constituting 13% (n=55/426), in contrast to 3% in both the GLASS-NL (n=3/98) and TCGA (n=6/223) datasets. This was also evident in WHO CNS5, due to the higher fraction of *CDKN2A/B* HD samples (CATNON: n=44/430, TCGA: n=15/252, GLASS-NL: n=8/98). The median age at diagnosis was significantly lower in the GLASS-NL (32 [18-70]) dataset in comparison to both CATNON (37 [18-82], p=5.92e-07) and TCGA (37 [14-74], p=9.95e-06). The observed differences in grade and age at diagnosis were presumably caused by differences in inclusion criteria.

### Continuous Grading Coefficient as a measure for grading/malignancy

For all datasets, methylation data were uploaded into the DNAm-based CNS tumor classifier for classification and copy number variation estimations. Three samples from the TCGA dataset exhibited 1p/19q codeletion (TCGA-CS-5394, TCGA-FG-7637, TCGA-VM-A8CA) and were therefore removed from all analyses (Supplementary Figure 1A).

The vast majority of the samples were classified as IDH-mutant astrocytoma (CATNON: 412/430, TCGA: 244/253, GLASS-NL-P: 93/98, GLASS-NL-R: 116/137). The highest fraction of A_IDH_HG samples was observed in the CATNON and GLASS-NL-R datasets (CATNON: 98/412, TCGA: 25/244, GLASS-NL-P: 4/93, GLASS-NL-R: 45/116) (Supplementary Figure 1A, Figure 1A). This aligns with the grade III inclusion criterion in CATNON and the higher proportion of progressed recurrent IDH-mutant astrocytoma in GLASS-NL (12).

**Figure 1:**
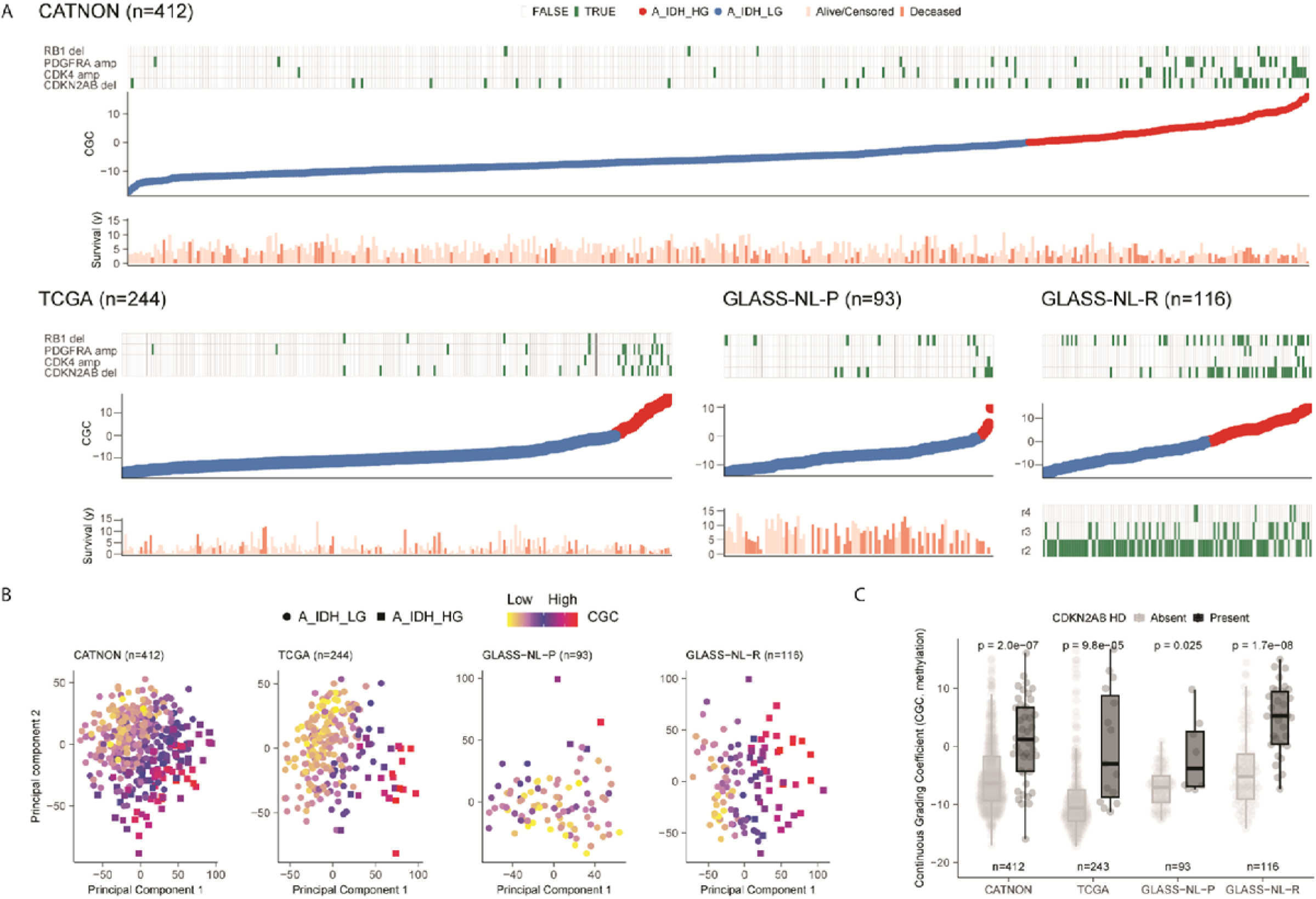
Continuous Grading Coefficient (CGC) as a tool to study malignancy in IDH-mutant astrocytoma. **A**, Samples of individual cohorts (CATNON, TCGA, GLASS-NL) ranked according to their CGC. As can be seen, specific copy number of events (RB1 HD, PDGFRA amplification, CDK4 amplification and CDKN2A/B HD), and overall survival were correlated with the CGC. Samples are colored based on astrocytoma subtype assignment (blue: A_IDH_LG, red: A_IDH_HG). **B,** Unsupervised PC1 and PC2 on the DNAm data showing spatial segregation of samples classified as IDH-mutant astrocytoma. **C,** Distribution of the CGC based on CDKN2A/B HD status across the different datasets. Samples with CDKN2A/B HD showed a significantly higher CGC. P-values determined by Wilcoxon signed-rank test.

Samples not classified as A_IDH_LG/A_IDH_HG were most often classified as oligodendroglioma (O_IDH, n=11) and oligosarcoma (OLIGOSARC_IDH, n=15) (Supplementary Figure 1A/B). Oligosarcoma is not recognized as a distinct tumor entity by the WHO CNS5, but represents oligodendroglioma with mixed oligodendroglial and sarcomatous morphology (27,28). Oligosarcoma forms a unique distinct methylation class that has been incorporated in the latest version (v12.8) of the CNS tumor classifier (28). Remarkably, samples classified as oligosarcoma did not harbor 1p/19q codeletion and had a lower tumor purity estimate compared to other IDH-mutant methylation subclasses (Supplementary Figure 1C, Supplementary Methods).

We utilized the calibrated classification probabilities derived from the CNS tumor classifier to generate a DNAm-based continuous grading coefficient (CGC). Formally, the CGC is calculated as the natural logarithm of the normalized classification probabilities between A_IDH_LG and A_IDH_HG. We then defined three revised astrocytoma subtype classes (low: CGC < −4.5, medium: CGC [-4.5-4.5], high: CGC > 4.5) based on the relationship between the CGC and the calibrated probability scores from the CNS tumor classifier in CATNON (Figure 2A). Clinical (age, sex, treatment), histological (necrosis and/or microvascular proliferation) and molecular (CNS tumor classifier class, CNV load and *CDKN2A/B* HD) characteristics of patients within each CGC class are presented in Supplementary Table 1. As may be expected, CGC class was positively associated with the number of samples being classified as WHO CNS5 grade 4 (CATNON: p<0.0001, GLASS-NL: p<0.0001, Fisher Exact Test, Figure 2B, Supplementary Table 1). However, a substantial proportion of WHO CNS5 grade 4 tumors were present in the CGC low and medium subgroups. Similarly, not all CGC-high tumors were WHO CNS5 grade 4. Our CGC groups were strongly associated with OS: median OS in CGC-low not reached (95% CI 8.2 – not reached), with CGC-medium 6.9 years (95% CI 5.7 – not reached) and CGC-high 3.4 years (95% CI 3.0 – not reached). All three CGC groups showed significantly different OS (CGC-medium vs CGC-low: HR 1.90, 95% CI [1.31-2.76]; p<0.001, CGC-medium vs CGC-high HR: 2.12 95% CI [1.33-3.36]; p=0.001). Multivariable Cox PH regression analysis on CATNON showed that the CGC groups were an independent prognostic factor (CGC-medium vs CGC-low: HR: 1.68, 95% CI [1.15-2.46] and CGC-high vs CGC-medium: HR: 3.42, 95% CI [2.12-5.51]) when adjusted for age, sex, *CDKN2A/B* HD, necrosis and/or microvascular proliferation and treatment with adjuvant/concurrent temozolomide (Figure 2D). Importantly, CGC cut-off values determined on CATNON also showed prognostic significance in TCGA (p<0.0001) and GLASS-NL-R (p<0.005) (Figure 2C). In these independent datasets, the CGC groups were significantly associated with CDKN2A/B HD (p<0.0001, Fisher Exact Test, Supplementary Table 1).

**Figure 2:**
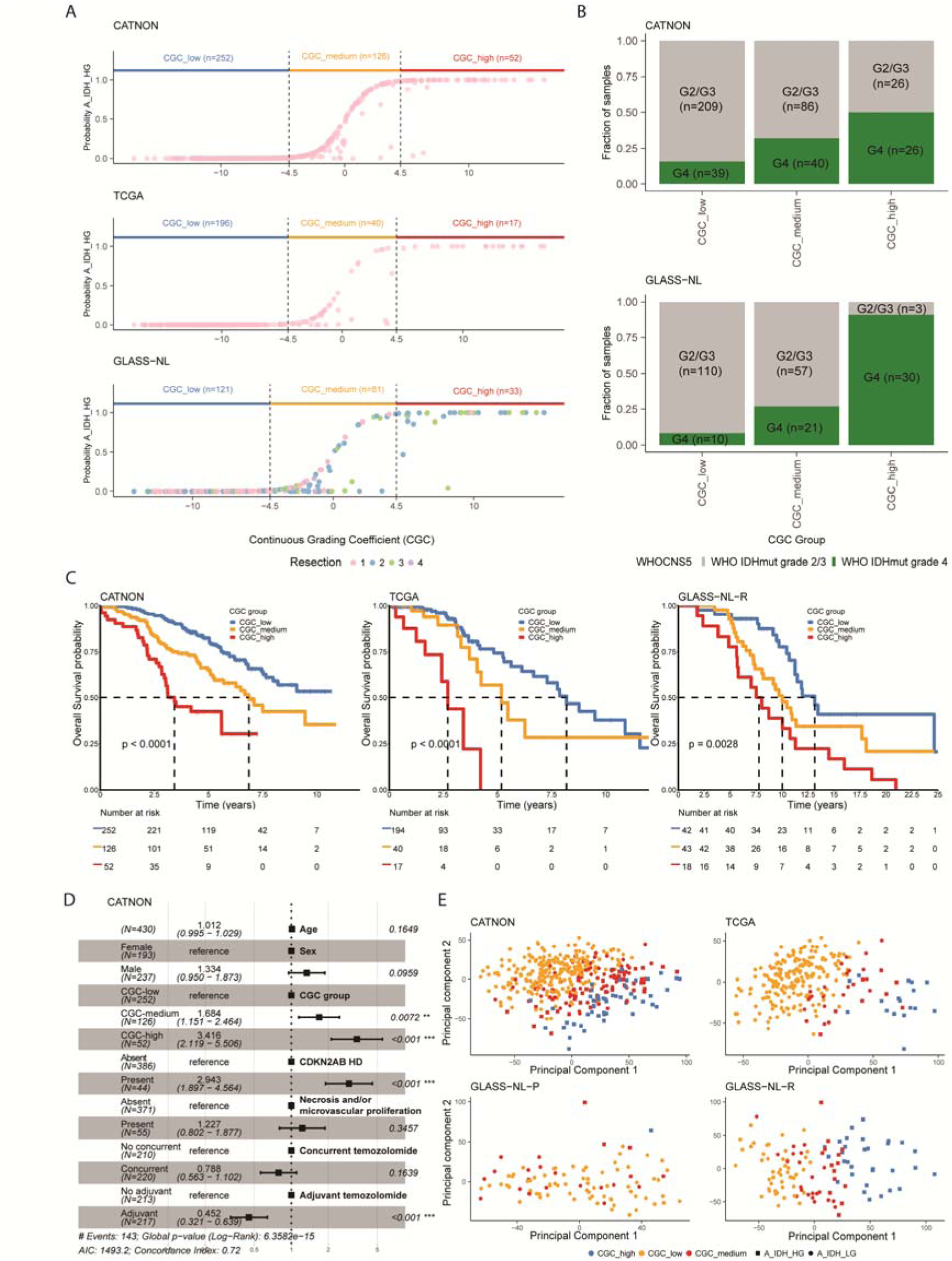
Evaluation of three CGC-based subgroups (low, medium, high) on CATNON, TCGA and GLASS-NL. **A,** Relationship between the CNS tumor classifier probability score and the CGC and resulting cut-off values for the three CGC groups (low: CGC < −4.5, medium: CGC [-4.5, 4.5], high: CGC > 4.5). **B,** Bar plot showing the fraction of samples according to WHO CNS5 within each CGC group. **C,** Kaplan-Meier overall survival curves stratified by CGC group. P-values were determined by log-rank test. **D,** Survival forest plots of predictive Cox proportional hazard models on the CGC groups corrected for WHO CNS5 criteria (*CDKN2A/B* HD and histological features) and treatment. **E,** Unsupervised PC1 and PC2 on the DNAm data showing spatial segregation of samples classified as IDH-mutant astrocytoma.

High CNV load is associated with worse outcome in IDH-mutant astrocytoma (7,13). We found that CGC croups were significantly associated with CNV load in all datasets (p<0.0001, Fisher Exact Test, Supplementary Table 1). CNV load was also associated with OS in a univariate analysis (CATNON: HR: 1.42 95% CI [1.02-1.97]; p=0.038, TCGA: HR: 2.42 95% CI [1.38-4.23]; p=0.0021, GLASS-NL-R: HR: 2.39 95% CI [1.45-3.96]; p<0.001).

We wondered to what extent our CGC captured the overall variability of DNAm profiles in our samples. To test this, we summarized the global DNAm profile of samples classified as IDH-mutant astrocytoma by PCA on the 10,000 most variable probes. Summarizing the DNAm profile by our three CGC groups captured the global methylation profile better compared to A_IDH_LG and A_IDH_HG (Figure 2E). However, we observed that the spatial distribution of methylation profiles along PC1 and PC2 was captured more effectively by the CGC for all datasets, emphasizing the efficacy of a continuous approach (Figure 1B).

High CGC values were not restricted to *CDKN2A/B* HD tumors alone (Figure 1A/C) and we next explored associations between the CGC and copy number events (Supplementary Figure 2). In addition to *CDKN2A/B* HD (CATNON: p=6.60e-07, TCGA: p=6.78e-05, GLASS-NL: p=1.88e-08), we also found associations with *SMARCA2* HD (CATNON: p=7.70e-05, TCGA: p=3.62e-04, GLASS-NL: p=6.68e-04), *RB1* HD (CATNON: p=6.40e-04, TCGA: p=0.033, GLASS-NL: p=0.019), *PTEN* HD (CATNON: p=2.73e-06, TCGA: p=0.12, GLASS-NL: p=1.69e-03), *CDK4* amplification (CATNON: p=5.15e-05, TCGA: p=0.016, GLASS-NL: p=0.046) and *PDGFRA* amplification (CATNON: p=5.15e-05, TCGA: p=0.019, GLASS-NL: p=0.024). It is interesting to note that the CGC captures more genetic events than the comparison between A_IDH_LG and A_IDH_HG tumors. For example, we identified a region on chromosome 10 encompassing the *PTEN* locus (CATNON: n=22; HR: 3.25 95% CI [1.86-5.66]; p<0.0001, GLASS-NL: n=8; HR: 1.42 95% CI [0.64-3.15]; p=0.39), that has thus far not been recognized as malignant marker in IDH-mutant astrocytomas.

### Global decrease in DNAm and hypermethylation of CpG islands associates with continuous grading coefficient

To elucidate what CpG sites associate with malignant transformation as defined by the CNS tumor classifier, we applied DMP analysis on the CGC using linear regression modelling for each dataset separately. In total, 8% of all tested probes were differentially methylated in the CATNON dataset and 8% in the TCGA dataset. In both datasets, the vast majority of differentially methylated probes were all hypomethylated in higher grade malignancies (CATNON: 99%, TCGA: 98%). Interestingly, although the vast majority of CpG sites had decreased methylation levels, those present on CpG islands were more often hypermethylated than hypomethylated (CATNON: p<2.2e-16, TCGA: p<2.2e-16, Figure 3A, Supplementary Table 2). Focusing on DMP present in both the 450k (TCGA) and EPIC (CATNON) array, we found that 61% of the hypermethylated probes from CATNON were shared in TCGA.

**Figure 3:**
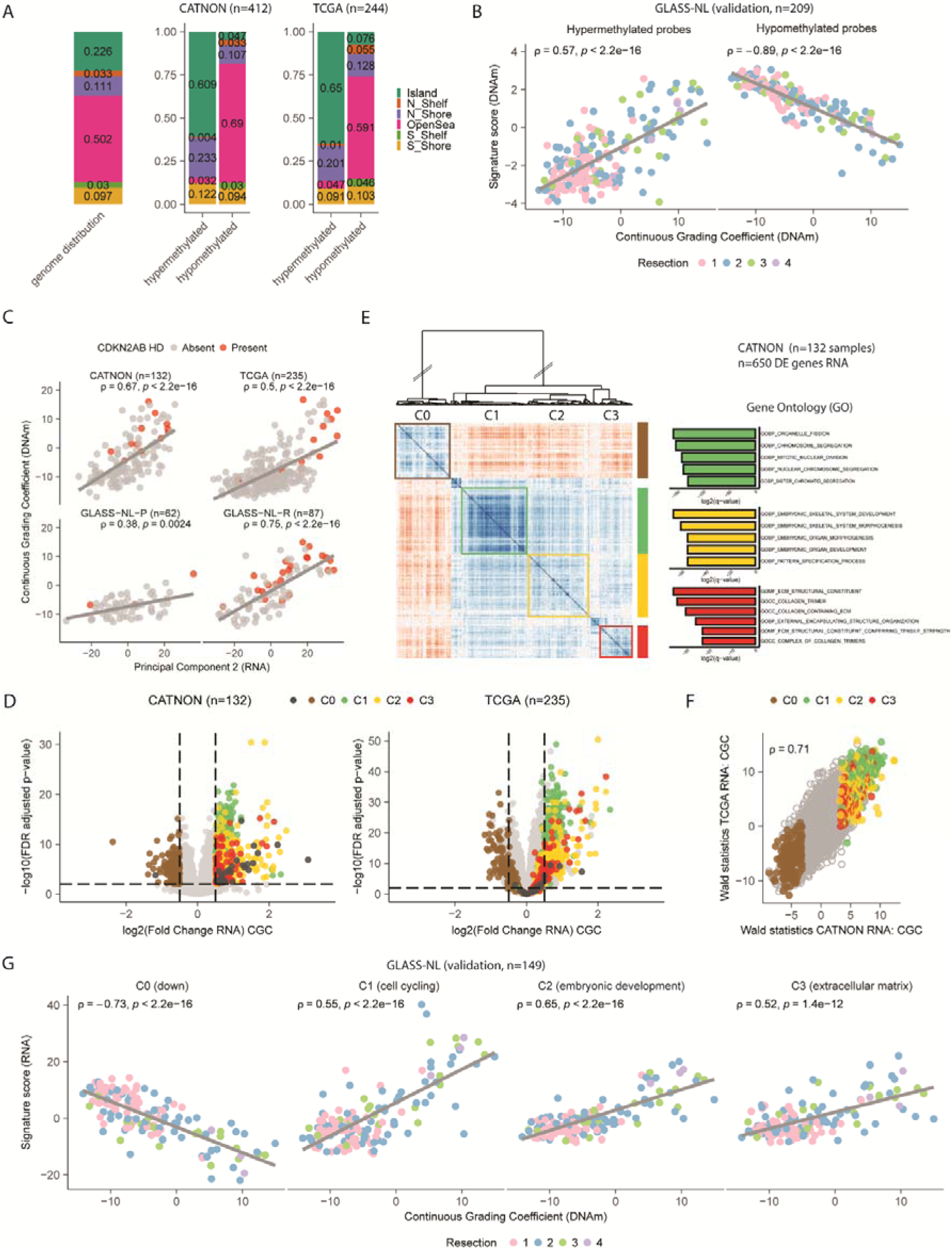
Supervised DNAm and RNA analysis on CATNON and TCGA with validation on the GLASS-NL dataset identifies gene clusters associated with malignancy. **A,** Distribution of probes belonging to CpG islands, shelfs (N_Shelf, S_Shelf), shores (N_Shore, N_Shelf) and the open sea across all hypomethylated and hypermethylated probes. Distributions are displayed separately for the DMPs resulting from independent analyses conducted on CATNON and TCGA. Genome-wide distribution of all probes is shown on the left as a reference. P-values were determined by Fisher’s Exact Test. **B,** Correlation between the DNAm-based signature scores of the hypermethylated and hypomethylated probes with the CGC in GLASS-NL. **C,** Spearman correlation between CGC and unsupervised PC2 of the transcriptomic data in all datasets. **D,** Volcano plots showing the per-gene RNA log2FoldChange on the CGC and the corresponding FDR adjusted p-value for both the CATNON and TCGA datasets. C0, C1, C2 and C3 genes are indicated. **E,** Recursive-based correlation plot on VST expression of the differentially expressed genes in CATNON. Three upregulated (C1: green, C2: yellow, C3: red) and one downregulated (C0: brown) cluster were distinguished. Gene Ontology enrichment resulted in significant hits for three clusters (C1: cell cycling, C2: embryonic development, C3: ECM). **F,** Correlation between the DGE tests on the CATNON and TCGA tests. For each gene the log2FoldChange divided by its standard error (Wald statistics) is indicated. **G,** Spearman correlation between the RNA-based signature scores for C0, C1, C2 and C3 and the CGC in GLASS-NL.

We subsequently validated the overlapping DMPs identified in the TCGA and CATNON datasets on GLASS-NL. The median M-value of the overlapping hypermethylated (n=149) and hypomethylated (n=12,708) probes both showed a correlation with the CGC and resection number in the GLASS-NL cohort (hypermethylated: ρ=0.57, hypomethylated: ρ=-0.89, Figure 3B). These correlations may be explained by an increased malignancy over time (12).

### Distinct transcriptional features are associated with our continuous grading coefficient

We included RNA-sequencing data of IDH-mutant 1p/19q non-codeleted CATNON and TCGA samples and findings were further validated on GLASS-NL-P and GLASS-NL-R. First, we conducted unsupervised PCA on the 1,000 most variably expressed genes for each dataset independently. In all datasets, the primary source of variation (PC1) was likely associated with tumor purity as demonstrated by the expression of neuronal genes, with neuron marker genes like *SLC12A5*, *TMEM130* and *SV2B* ranking among the top 50 contributors (24). PC2 was however strongly associated with the DNAm-based CGC (CATNON: ρ=0.67, TCGA: ρ=0.50, GLASS-NL-P: ρ=0.38, GLASS-NL-R: ρ=0.75, Figure 3C).

Gene-level differential regression analysis on samples from the CATNON and TCGA was then performed to find genes associated to our substantiating the CGC. To ensure we are mainly investigating tumor-intrinsic signal, we estimated tumor purity and investigated its influence on the outcome of the DGR analysis on the CGC (Supplementary Figure 3, Supplementary Methods). We did not observe a clear dependency of tumor purity on the outcome of the DGR analysis (CATNON: ρ=-0.23, TCGA: ρ=0.28) nor a difference in tumor purity based on *CDKN2A/B* HD status (CATNON: p=0.45, TCGA: p=0.39). We therefore did not perform additional purity corrections on our datasets.

DGR on the CGC revealed a total of 650 differentially expressed genes in the CATNON dataset (Supplementary Table 2). The differential gene expression was skewed, with a higher proportion (77%) of genes showing increased expression in samples with a higher CGC (Figure 3D). We then performed recursive-based correlation clustering (18) and identified four distinct gene clusters (Figure 3E, C0/C1/C2/C3). The clusters that showed increased expression in more malignant samples (high CGC) were associated with specific biological functions: C1 (cell cycling, n=175), C2 (embryonic development, n=176) and C3 (ECM, n=97) (Figure 3E). C0 (n=149), which contained genes with a decreased expression in more malignant samples, could not be attributed to a biological function. DGR analysis on the TCGA dataset resulted in 667 differentially expressed genes (Figure 3D), with a strong overlap with those identified with the CATNON analysis (58%). Also, the majority of the differentially expressed genes were upregulated (72%). Despite the differences in study designs, we observed a strong correlation across both datasets in outcome of the tests performed (ρ=0.71, Figure 3F).

Subsequently, we further examined the genes present within each of the transcriptional clusters. C1 contained genes associated with histones (e.g. *H3C2*, *H2BC9*, *H2BC11*), transcription factors (e.g. *E2F1*, *E2F8*, *FOXM1*), kinesins (e.g., *KIF14*, *KIFC1*), DNA replication (e.g. *RAD51*, *EXO1*, *CENPK*) and cell cycle control (e.g. *CHEK1*, *CCNB1/2*, *CDK6*). Of note, established markers for cell cycling, such as *MKI67* and *TOP2A* were found. C2 contained developmental transcription factor genes, such as members of the *HOX* (n=18), *PAX* (n=3) and *TBX* (n=2) gene family. Within C3, genes associated with ECM formation were found, such as *COL1A1* and *COL1A2*. On the contrary, C0 comprised many long non-coding RNA genes (n=81). We compared expression levels of C0 genes among purified human CNS cell types (Supplementary Methods, Supplementary Figure 4), and observed high expression levels in mature astrocytes as compared to fetal astrocytes (p<0.001). Conversely, upregulated genes (C1-C3) were higher expressed in fetal astrocytes (p<0.001). These combined findings suggest tumor dedifferentiation during malignant transformation.

PCA was conducted individually for each cluster (C0-C3) to derive a per tumor-sample representative value. This value summarized the expression pattern of genes within the respective cluster, as represented by PC1. Importantly, the differentially expressed genes used to define these RNA signatures were identified based on results from CATNON alone. When these values were calculated on the GLASS-NL dataset, we found correlations between the CGC and all RNA signature scores (C0: ρ=-0.73, C1: ρ=0.55, C2: ρ=0.65 and C3: ρ=0.52, Figure 3G).

We utilized our snRNA-seq dataset (vanHijfte_2023), which included six matched primary and recurrent IDH-mutant astrocytoma samples from three patients (Figure 4A). For all patients, we selected the primary samples to be A_IDH_LG and the recurrent A_IDH_HG. Uniform Manifold Approximation and Projection (UMAP) showed distinct cell types and the previously reported tumor cell states (astro-like, oligo-like and stem-like, Figure 3A, Supplementary Figure 5A). Interestingly, at tumor recurrence/high-grade, the oligo-like and astro-like cell states were less pronounced and a substantial tumor cell cluster could not reliably be defined using our marker gene set. We then overlaid our transcriptional signatures (C0-C3) with this dataset and found that C0, C1 and C2 were predominantly expressed in tumor cells (Figure 4B/C). More specifically, C0 and C1 showed high expression in astro-like and proliferating tumor cells respectively (25,29). Genes from C3 were equally expressed by endothelial cells/pericytes and undetermined tumor cells (Figure 4B/C). In line with our RNA-sequencing results, expression of C0 genes decreased and C1/C2/C3 increased during malignant transformation. In summary, our data suggest that malignant transformation of IDH-mutant astrocytoma is associated with an upregulation of cell cycling, embryonic development and ECM genes. This was accompanied by decreased expression of astro-like state genes.

**Figure 4:**
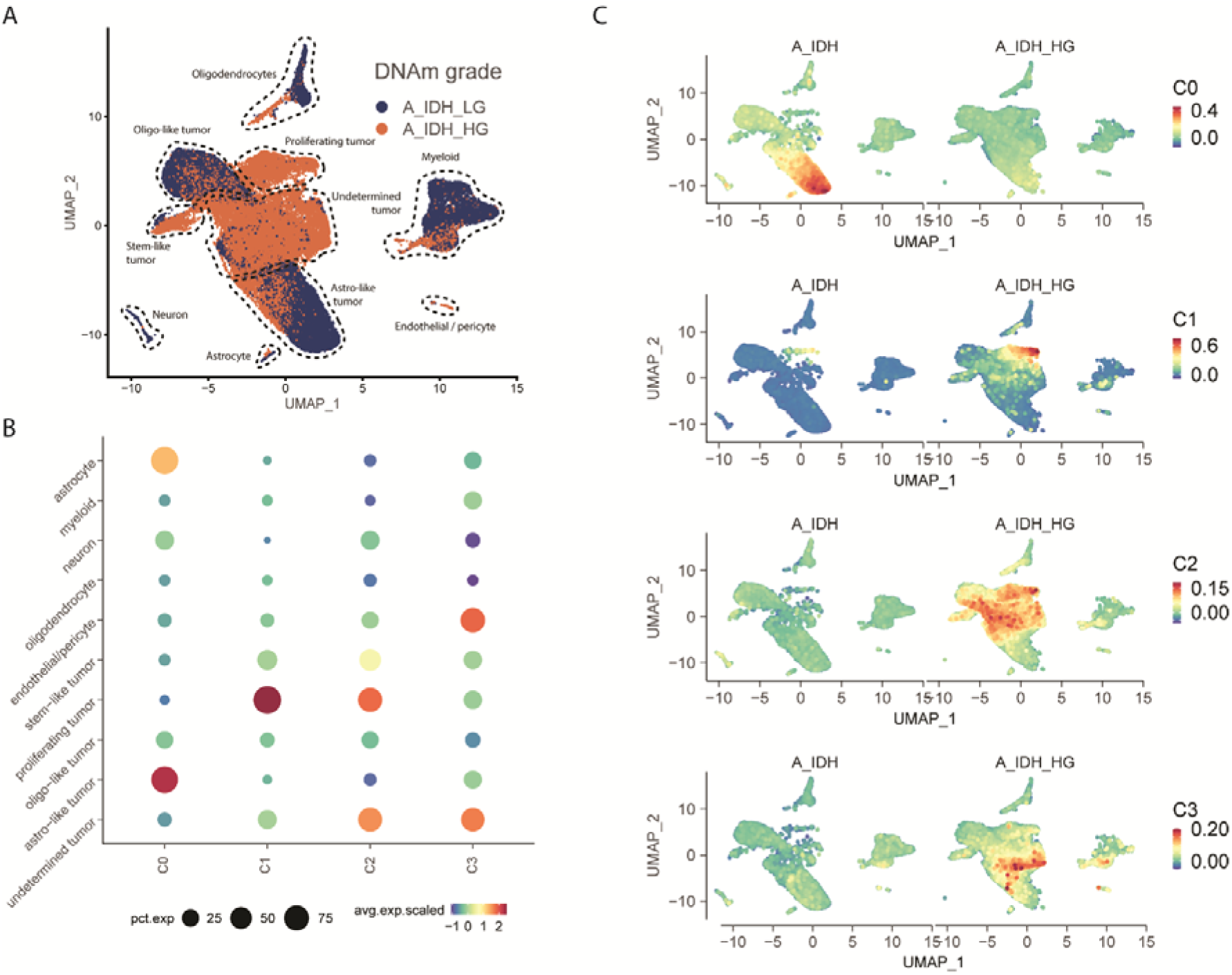
Single-nucleus RNA-sequencing (A_IDH_LG: n=3, A_IDH_HG: n=3) validates bulk DGE results and shows enrichment of gene clusters in select cell subpopulations. **A,** UMAP projection showing cell-type annotations for all tumors combined. **B,** Dot plot showing the enrichment score of each of the bulk RNA-sequencing clusters (C0, C1, C2, C3) identified in our DGR analysis. **C,** UMAP projection showing enrichment scores of the downregulated (C0), cell cycling (C1), embryonic development (C2) and ECM (C3) clusters for A_IDH_LG and A_IDH_HG separately.

### Hypermethylation and upregulation of embryonic development genes in more malignant IDH-mutant astrocytoma

We next integrated the methylome differences with transcriptional changes of the corresponding genes (Figure 5A). The differentially hypermethylated probes (n=149) were associated with 63 genes from our RNA expression data (Supplementary Table 2). Pathway enrichment analysis on these hypermethylated genes revealed significant hits for high-CpG-density promoter (HCP) genes bearing the histone H3 trimethylation mark at K27 (H3K27Me3) in brain (FDR-adjusted p-value=4.8e-14).

**Figure 5:**
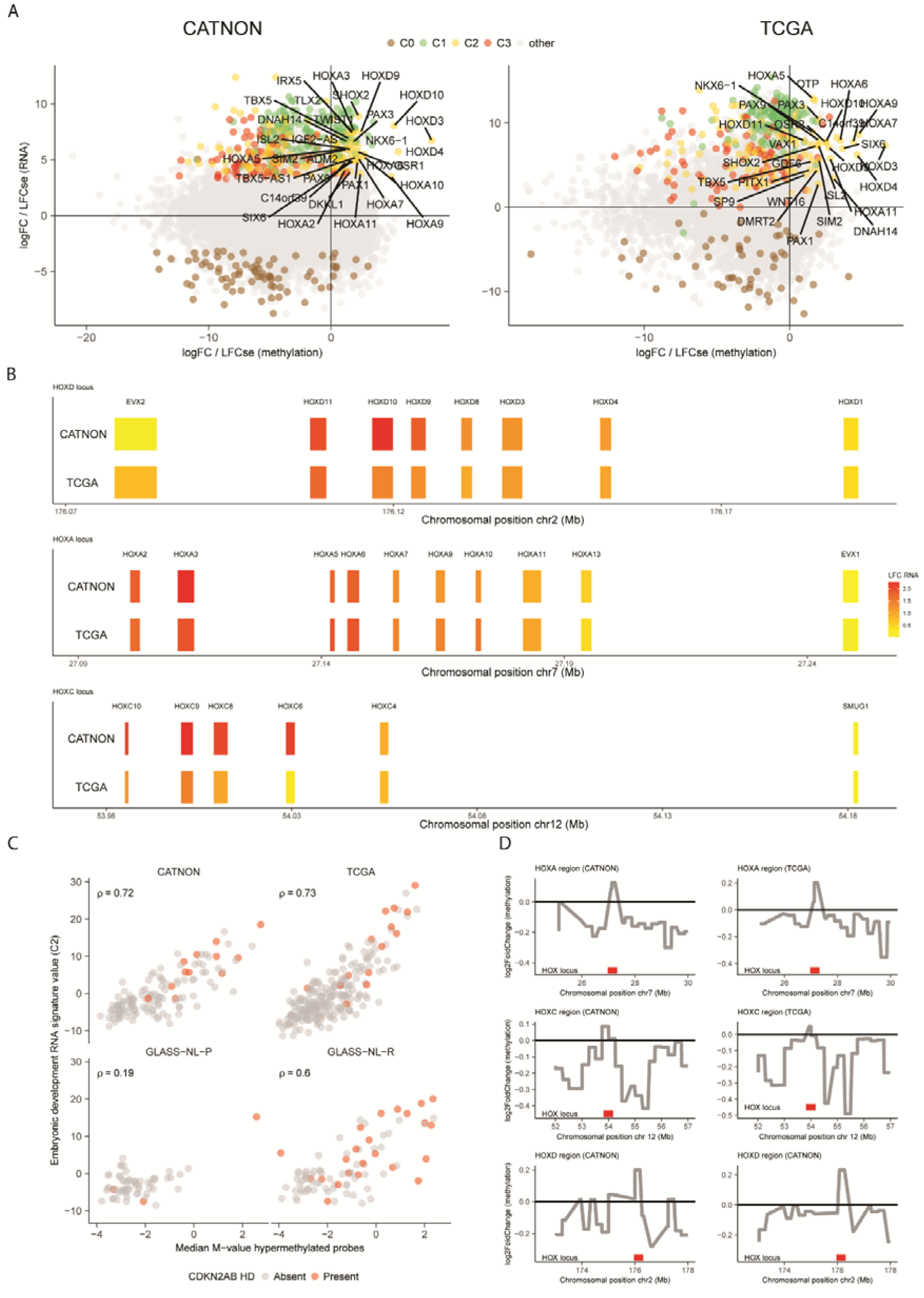
Supervised analysis shows co-existence of upregulated and hypermethylated embryonic development genes. **A,** Correlation between the DMP (x-axis) and DGR (y-axis) across the CATNON and TCGA datasets. **B,** Log2FoldChange of the CGC on RNA VST expression for the *HOXD* (chr2), *HOXA* (chr7) and *HOXC* (chr12) loci. Chromosomal positions are indicated in megabases (Mb). **C,** Correlation between the embryonic development RNA signature scores (C2) and the median M-value of the hypermethylated probes. *CDKN2A/B* HD status is indicated in red. **D,** Log2FoldChange of the CGC on the DNAm M-values for the *HOX* loci (*HOXA, HOXC, HOXD)* and the surrounding regions on the CATNON and TCGA datasets.

Comparison of the hypermethylated genes with transcriptional changes of the corresponding genes revealed a strong overlap with the upregulated embryonic development cluster (C2, n=27/63 genes). Thus, although the vast majority of the genes were hypomethylated with increased malignancy/CGC, we found hypermethylated CpGs to be associated with genes that were, paradoxically to what may be expected, transcriptionally upregulated. These hypermethylated and upregulated C2 genes encode for key developmental transcription factor genes, such as members of the *HOX* (n=11), *PAX* (n=2) and *TBX* (n=1) family of genes.

The per-sample median M-value of the hypermethylated probes correlated strongly with the C2 signature score on both the CATNON (ρ=0.72) and TCGA (ρ=0.73) datasets (Figure 5C). We also find a modest correlation in GLASS-NL-P (ρ=0.19), which may be expected due to the low number of high-grade primary samples. Conversely, a strong correlation was evident GLASS-NL-R (ρ=0.60) (Figure 5C).

### Locus-specific hypermethylation and upregulation of HOX genes in tumor cells

When we further examined the expression of the *HOX* genes of C2, we observed gene-ordered differential expression along the different gene clusters, where genes positioned at the 5’ end of the loci show a higher expression with higher CGC scores and gradually decrease downstream in both CATNON and TCGA (Figure 5B). This suggests coordinated derepression of enhancer sequences up/downstream of the gene cluster. Indeed, regions surrounding the *HOX* loci indeed were hypomethylated, supporting the hypothesis of derepression of enhancers (Figure 5D). Importantly, in both the bulk and single-cell data increased *HOX* expression of those present in C2 was not restricted to tumors with *CDKN2A/B* HD (Supplementary Figure 5B).

### Histopathological features that associate with continuous grading coefficient

We investigated the correlation between thirteen histological features and the CGC through a multivariate linear model to gain insight into their association with tumor malignancy. For the CATNON dataset, thirteen histological features were scored by a panel of seven international neuropathologists: cell density, calcifications, increased number of blood vessels, microvascular proliferation, neoplastic appearing astrocytes and oligodendrocytes, giant cells, (miniature) gemistocytes, nuclear pleomorphism, microcysts and mucoid degeneration, necrosis and mitotic count (30). Of these, the mitotic index (p=0.0093) and giant cells (p=0.042) were significantly associated with the CGC. This is in line with earlier work, which showed prognostic significance of the mitotic index (30). We next correlated these variables with our RNA signatures and found that samples with a high mitotic index (>2 mitoses per 10 40x consecutive high-power fields) and presence of giant cells had a higher C1 cell cycling (mitotic index: p=0.015, giant cells: p=0.029) and C2 embryonic development (mitotic index: p=0.003, giant cells: p=0.0019) signature value. Our data revealed that increased cell cycling at the gene level is correlated with an elevated mitotic index.

### Hypermethylation phenotype and high expression of embryonic development genes is associated with poor survival

We categorized samples into low and high-risk groups for the C2 and hypermethylation signatures individually, based on the sign (negative or positive) of the C2 signature and PC1 associated with the M-value of hypermethylated probes (n=149), respectively. Both high RNA expression of C2 genes (p<0.0001, HR: 3.90 95% CI [2.13-7.13]) and the hypermethylation phenotype (p<0.0001, HR: 2.42 95% CI [1.73-3.38]) were significantly associated with shorter survival in a univariate analysis on the CATNON dataset (Figure 6A/B). These associations remained present after adjusting for age, sex, *CDKN2A/B* HD, microvascular proliferation/necrosis and adjuvant/concurrent temozolomide using a multivariable Cox PH model (C2 signature: p=0.014, HR: 2.42 95% CI [1.20-4.87], hypermethylation phenotype: p=0.0048, HR: 1.73 95% CI [1.18-2.52], Figure 6C/D). These molecular signatures hold more prognostic value compared to the mitotic index (p=0.053, HR: 1.66 [0.99-2.77]) in a univariate analysis.

**Figure 6:**
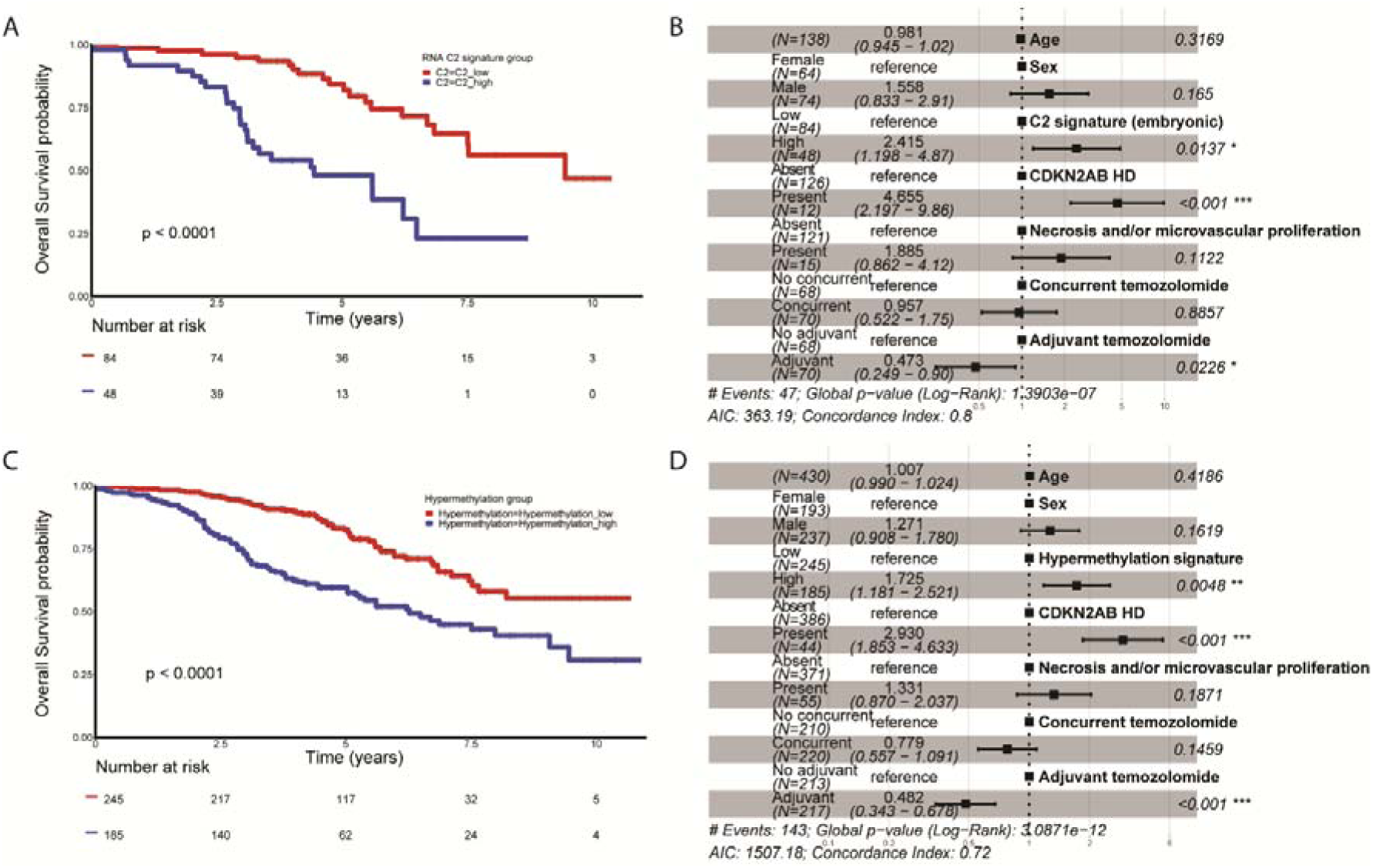
Survival analysis of RNA and methylation signatures using the CATNON dataset. **A,C** Kaplan-Meier overall survival curves of CATNON stratified by RNA C2 signature (A) and hypermethylation phenotype (C) risk groups. **B/D**, Survival forest plots of predictive Cox proportional hazard models on the C2 signature (B) and hypermethylation phenotype (D) corrected for age, sex, *CDKN2A/B* HD, treatment and histology.

The C2 RNA signature (p<0.0001, HR: 3.11 [1.7-5.6, 95% CI] and hypermethylation phenotype (p<0.0001, HR: 3.43 [1.93-6.11, 95% CI]) were also associated with shorter survival on the TCGA dataset (Supplementary Figure 6). The signature values at recurrence on GLASS-NL also showed prognostic significance (C2 RNA: p=0.0047, HR: 2.58 [1.30-5.11, 95% CI], hypermethylation: p=0.012, HR: 1.92 [1.15-3.23, 95% CI], Supplementary Figure 6).

## Discussion

In this study, we aimed to enhance the understanding of malignant transformation in IDH-mutant astrocytomas at a multi-domain molecular level. To do so, we generated RNA-sequencing data from IDH-mutant astrocytomas included in the CATNON randomized phase III clinical trial and combined it with additional multi-domain high-throughput data, along with multi-omics data from GLASS-NL and TCGA. We defined cut-off values for the methylation-based CGC for improved prognostication and found prognostic (epi)genetic and transcriptional markers that converge on three pathways: upregulation of cell-cycling genes, modification of the ECM, and tumor cell dedifferentiation (both by a reduced expression of mature astrocyte genes and upregulation of developmental genes). Importantly, altered expression of these three signatures was not restricted to tumors with *CDKN2A/B* HD, which demonstrates the additional prognostic power of our analyses.

In recent years, the diagnostic classification of primary brain tumors has shifted from primarily histopathology to a more precision-oriented approach that incorporates molecular markers to optimize prognostication. We developed an objective continuous grading coefficient, based on a DNAm-based classifier to assign a continuous grade per patient (5). We defined three astrocytoma groups based on the CGC, which adds an intermediate risk group. These subclasses showed distinct OS, correlated with WHO CNS5 and high CNV load. Prognostic significance of these subgroups was further confirmed in our multivariable COX regression analysis where they emerged as independent prognostic factor.

In all datasets, the CGC correlated strongly with the molecular marker *CDKN2A/B*, which is already incorporated in the WHO CNS5 classification. We also found members of the RB1 pathway to be individually associated with the CGC, including *CDK4* amplification and *RB1* HD. Additionally, we identified *PDGFRA* amplification to be associated with the CGC. These genes were described to correlate with malignancy in earlier work (31,32). This supports the CGCs role as generic grading marker, since we used it to identify these associations, and it therefore encapsulates information on all these individual malignant genetic events. Also, continuity of tumor grading was supported since the incidence of these malignant events correlated with the value of the CGC. Additionally, deletion of the locus encompassing *PTEN* was found to be associated with both the CGC and OS in the CATNON dataset. This alteration has not been described as established molecular marker related to patient outcome in IDH-mutant astrocytomas (32).

After deducing transcriptional profiles associated with the CGC, we found multiple genes from the *HOX* gene clusters among our upregulated embryonic development genes. These genes exhibited a spatial sequential expression pattern reminiscent of that observed during embryogenesis. Despite genome-wide DNA-demethylation at progression, the *HOX* loci showed an increase in methylation levels and an increased RNA-expression. Hypermethylation of *HOX* genes has been described previously in glioma (33–36). However, there is no consensus regarding the subsequent epigenetic silencing or reactivation of these genes (33,35). We showed transcriptional upregulation and hypermethylation of the *HOXA*, *HOXC* and *HOXD* gene clusters in higher-grade IDH-mutant astrocytomas across multiple datasets. The ordered differential expression of *HOX* genes accompanied by focal and targeted hypermethylation, particularly located near the *HOX* genes, raise intriguing questions about their potential significance in malignant transformation. Biologically, hypermethylation of genes is often associated with gene repression (37). Regulatory elements of *HOX* genes reside in topologically associating domains (TADs), which are involved in long-distance gene regulation (38). In IDH-mutant gliomas, it has been shown that CTCF-binding is disrupted, leading to a loss of insulation between TADs and aberrant gene expression (39). Derepression of long-distance regulatory elements might counterbalance the repressive effect on promotor hypermethylation. In IDHwt glioblastoma, coexistence of hypermethylation and upregulation of *HOX* genes was linked H3K27me23 depletion and the use of alternative promotors (34). Future experimental work on IDH-mutant glioma is needed to elucidate the underlying mechanism.

Our analyses elucidated the role of tumor dedifferentiation in shaping malignant transformation of IDH-mutant astrocytomas. We observed downregulation of genes associated with mature astrocytes. Concurrently, we found an upregulation of developmental genes, indicative of a less differentiated state. We also revealed increased proliferation and higher expression of ECM remodeling genes to be associated with malignancy.

While histological grading, based on traditional morphological assessments, has long been a cornerstone in tumor classification, our study underscores the superior prognostic efficacy of molecular signatures. The clinical implication of our CGC is that it allows an objective means to grade IDH-mutant gliomas, with tumor grade being used as an aid in clinical decision making. Although we argue that grade is a continuum, the cut-off points can identify the most indolent and the most aggressive tumors more accurate than grading based on WHO CNS5 and the CNS tumor classifier subgroups.

## Author’s Contributions

**S.A. Ghisai:** Conceptualization, Methodology, Formal Analysis, Validation, Visualization, Data curation, Writing – review & editing, Writing – original draft, **L. van Hijfte**: Formal Analysis, Validation, Visualization, Investigation, Data curation, Resources, Writing – review & editing, **W.R. Vallentgoed**: Conceptualization, Methodology, Formal Analysis, Validation, Investigation, Data curation, Resources, Writing – review & editing, **C.M.S. Tesileanu**: Methodology, Investigation, Data curation, Resources, Writing – review & editing, **I. de Heer**: Investigation, Data curation, Writing – review & editing, **J.M. Kros**: Investigation, Resources, Writing – review & editing, **M. Sanson**: Resources, Writing – review & editing, **T. Gorlia**: Resources, Writing – review & editing, **W.Wick**: Resources, Writing – review & editing, **M.A. Vogelbaum**: Resources, Writing – review & editing, **A.A. Brandes**: Resources, Writing – review & editing, **E. Franceschi**: Resources, Writing – review & editing, **P.M. Clement**: Resources, Writing – review & editing, **A.K. Nowak**: Resources, Writing – review & editing, **V. Golfinopoulos**: Writing – review & editing, Data curation, Supervision, **M.J. van den Bent**: Conceptualization, Methodology, Funding acquisition, Writing – original draft, Writing – review & editing, Project administration, Supervision, **P.J. French**: Conceptualization, Methodology, Funding acquisition, Writing – original draft, Writing – review & editing, Project administration, Supervision, **Y. Hoogstrate**: Conceptualization, Methodology, Data curation, Writing – original draft, Writing – review & editing, Project administration, Supervision

## Acknowledgements

The CATNON study was funded by Merck Sharp & Dohme (formerly Schering-Plough) through an educational grant and provision of temozolomide. The RNA-sequencing was supported by grants from by the Vereniging Heino ‘Strijd van Salland’ and by Stichting STOPhersentumoren. The further molecular studies were supported by grant GN-000577 from The Brain Tumour Charity and grant 10685 from the Dutch Cancer Society. The translational research was further supported by the NRG (grants U10CA180868 and U10CA180822), Cancer Research UK grant CRUK/07/028, and Cancer Australia (project grants 1026842 and 1078655). We thank the patients and their relatives for their willingness to participate to the CATNON study. We also thank all sites and their staff for contributing to the CATNON study. The CATNON study protocol was prepared and the study database was developed, housed, and analysed by the EORTC. We acknowledge the support of the CATNON study by the staff at the EORTC Headquarters in Brussels, Belgium, the NRG Oncology (formerly the Radiation Therapy Oncology Group) staff at the American College of Radiology; the staff at the Australian National Health and Medical Research Council (NHMRC) Clinical Trials Centre (COGNO Coordinating Centre); and the staff at MRC Clinical Trials Unit, London UK.

---

**Supplementary Figure 1:**
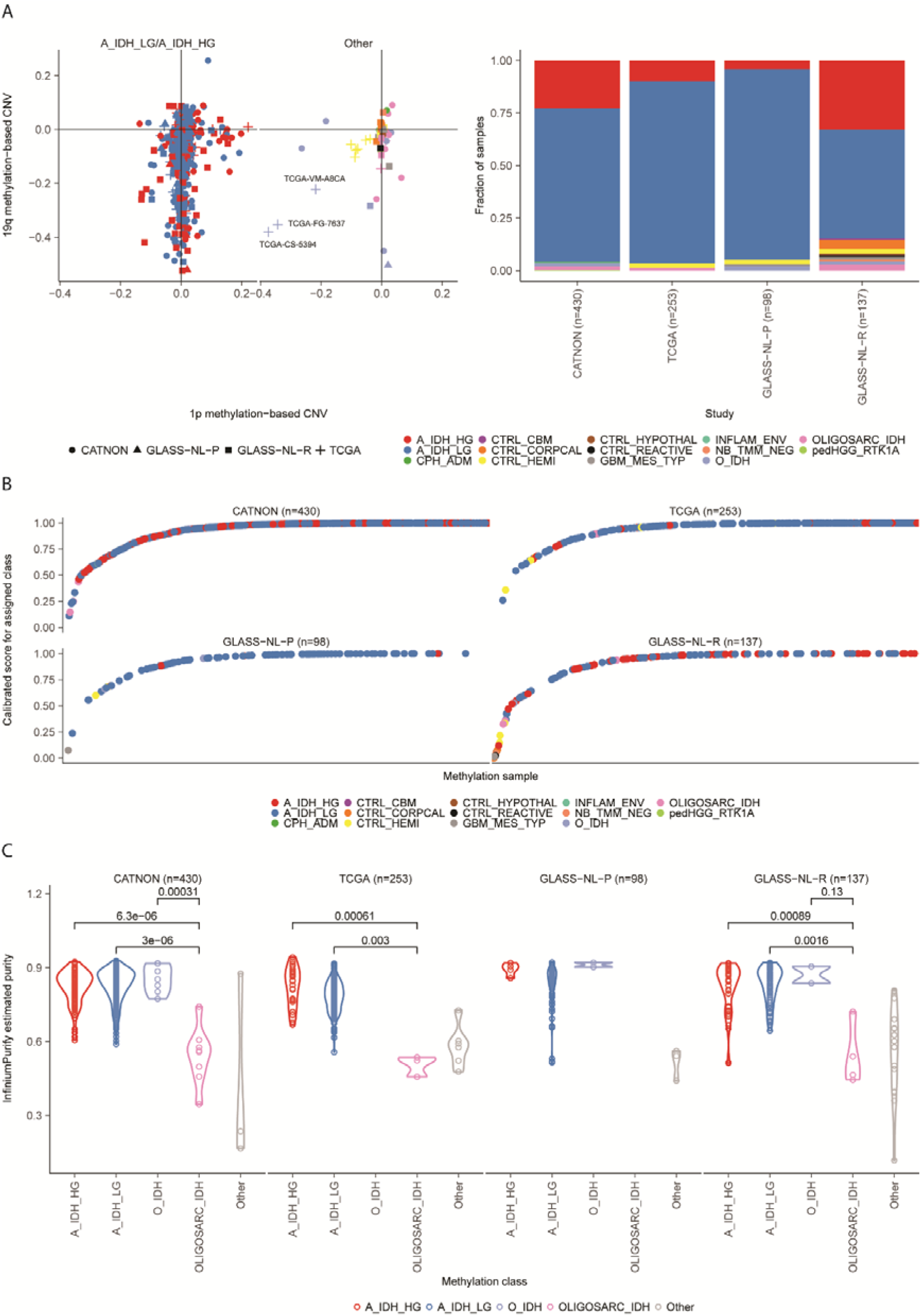
Assignment of CNS tumor subtypes by the CNS tumor classifier and assessment of tumor purity. **A,** Methylation-based copy number variation estimates for chr1p and chr19q (left) and subtype classification (right) for the CATNON, TCGA, GLASS-NL-P and GLASS-NL-R datasets. **B,** Calibrated scores for the assigned subtype classes per dataset. **C,** Estimated purities for the assigned methylation classes. P-values determined by Wilcoxon signed-rank test.

**Supplementary Figure 2:**
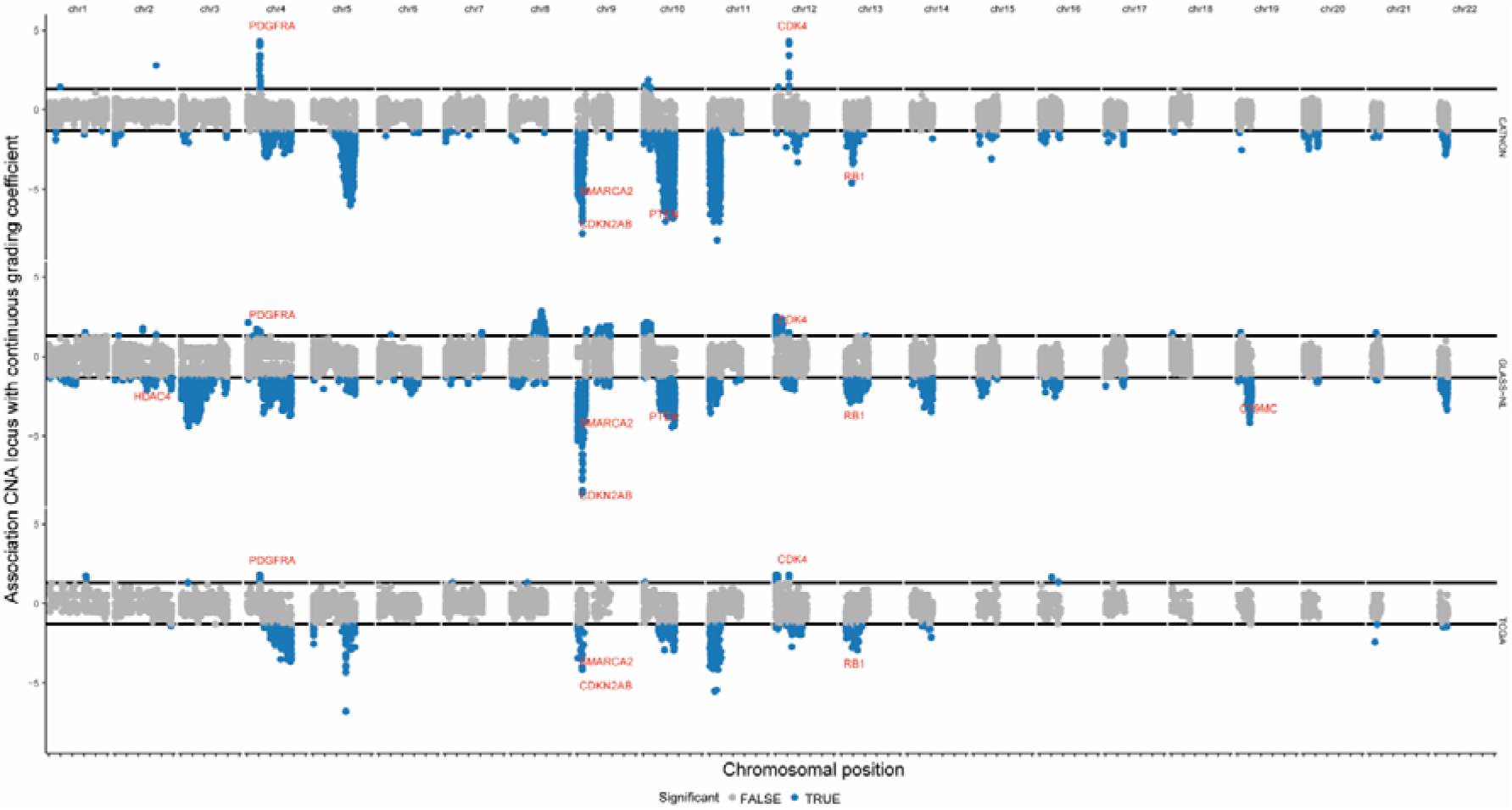
Associations (log10(FDR-adjusted p-value)) between the CNA locus and the Continuous Grading Coefficient (CGC) for the CATNON, TCGA and GLASS-NL datasets. P-values were determined by the Wilcoxon signed-rank test comparing per genomic bin the CGC between samples with and without a deletion or gain event within the respective bin.

**Supplementary Figure 3:**
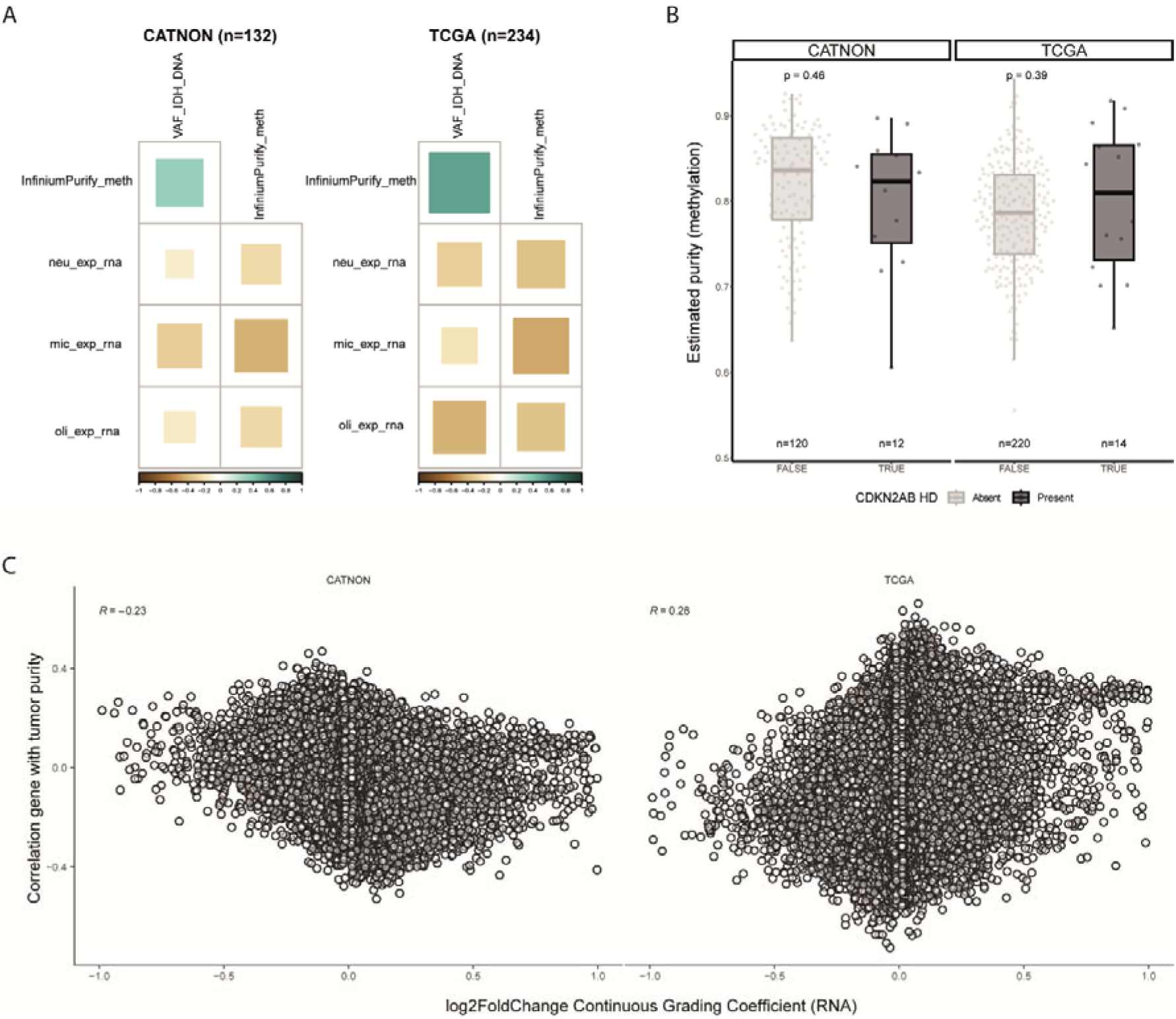
Association between tumor purity and outcome of the DGR analysis. **A,** Correlation between the expression of oligodendrocyte, microglia and neuron expression marker genes (24) and tumor purity estimation methods (VAF *IDH* mutation, InfiniumPurify) for both the CATNON and TCGA datasets. **B,** Distribution of InfiniumPurify estimated purities based on the presence of *CDKN2A/B* HD. For both the CATNON and TCGA dataset no significant difference in tumor purity was observed. P-values determined by Wilcoxon signed-rank test. **C,** Correlation of the log2FoldChange (RNA) of the Continuous Grading Coefficient (CGC) with InfiniumPurify estimated purities.

**Supplementary Figure 4:**
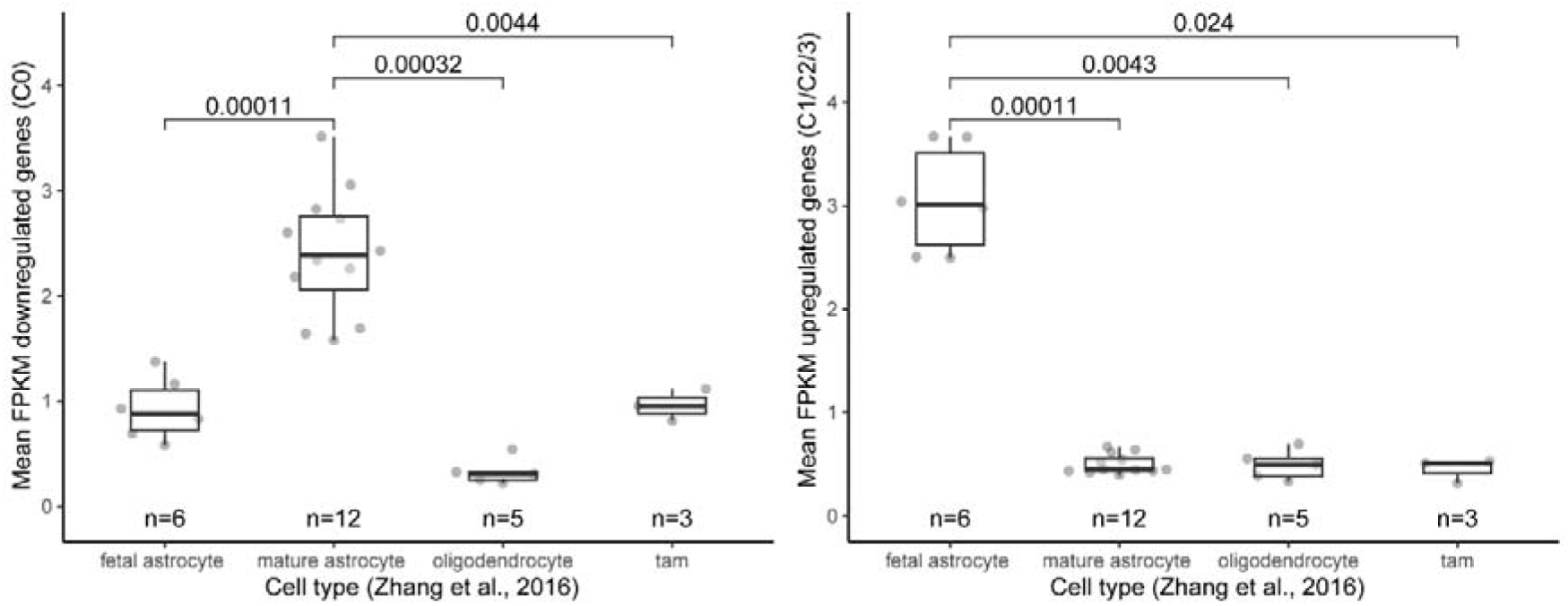
Mean FPKM of downregulated (C0, left) and upregulated (C1/C2/C3) transcriptional cluster genes identified in our DGR analysis across fetal astrocytes (n=6), mature astrocytes, oligodendrocytes and tumor-associated macrophages (40). P-values determined by Wilcoxon signed-rank test.

**Supplementary Figure 5:**
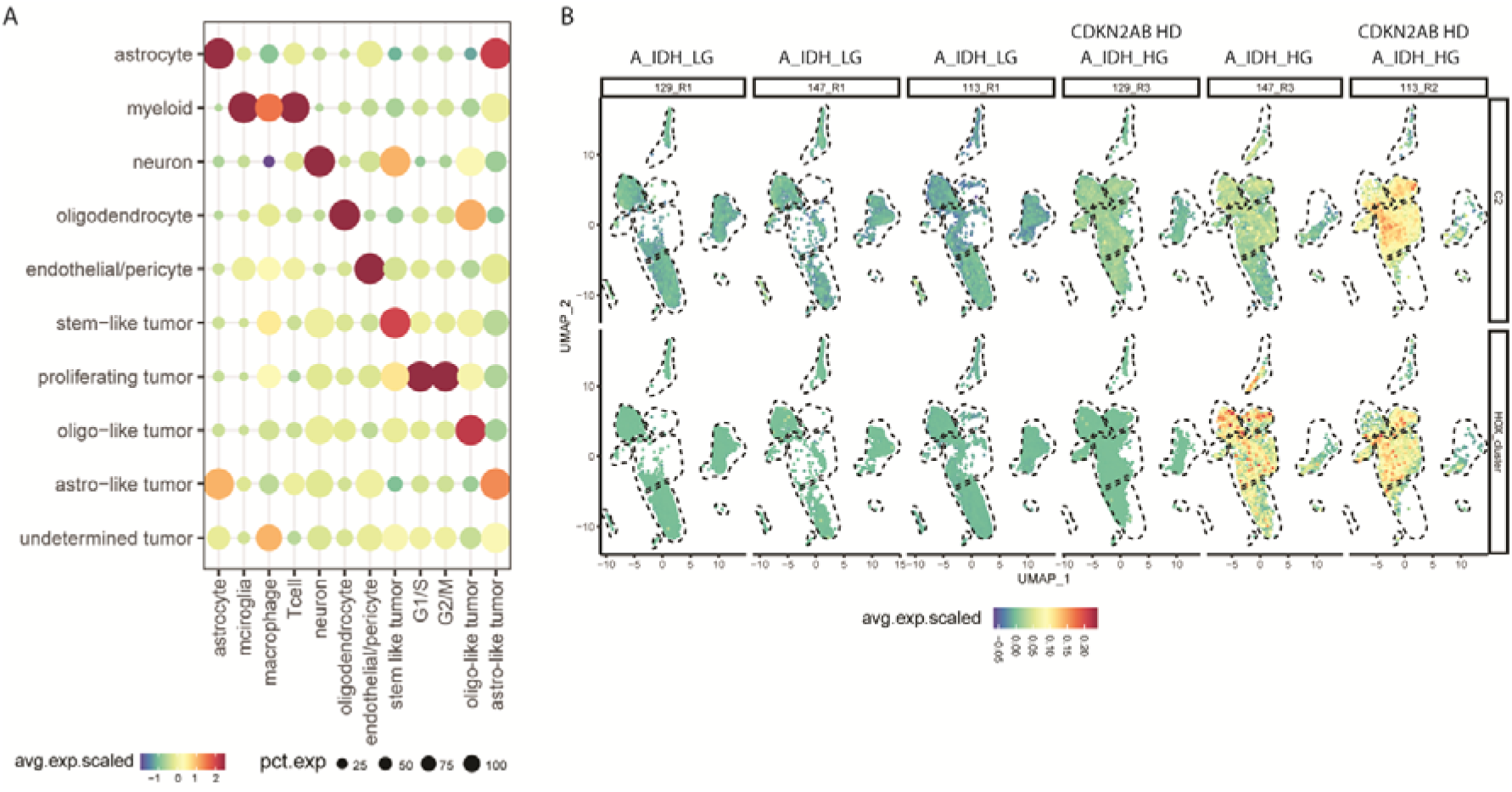
Single-nucleus RNA sequencing enrichment scores on methylation-based low-grade and high-grade samples. **A,** Dot plot showing the enrichment scores of marker genes for each of the assigned cell types. **B,** UMAP projections showing enrichment scores of the embryonic development (C2) cluster and *HOX* genes (from C2). *CDKN2A/B* HD status is indicated.

**Supplementary Figure 6:**
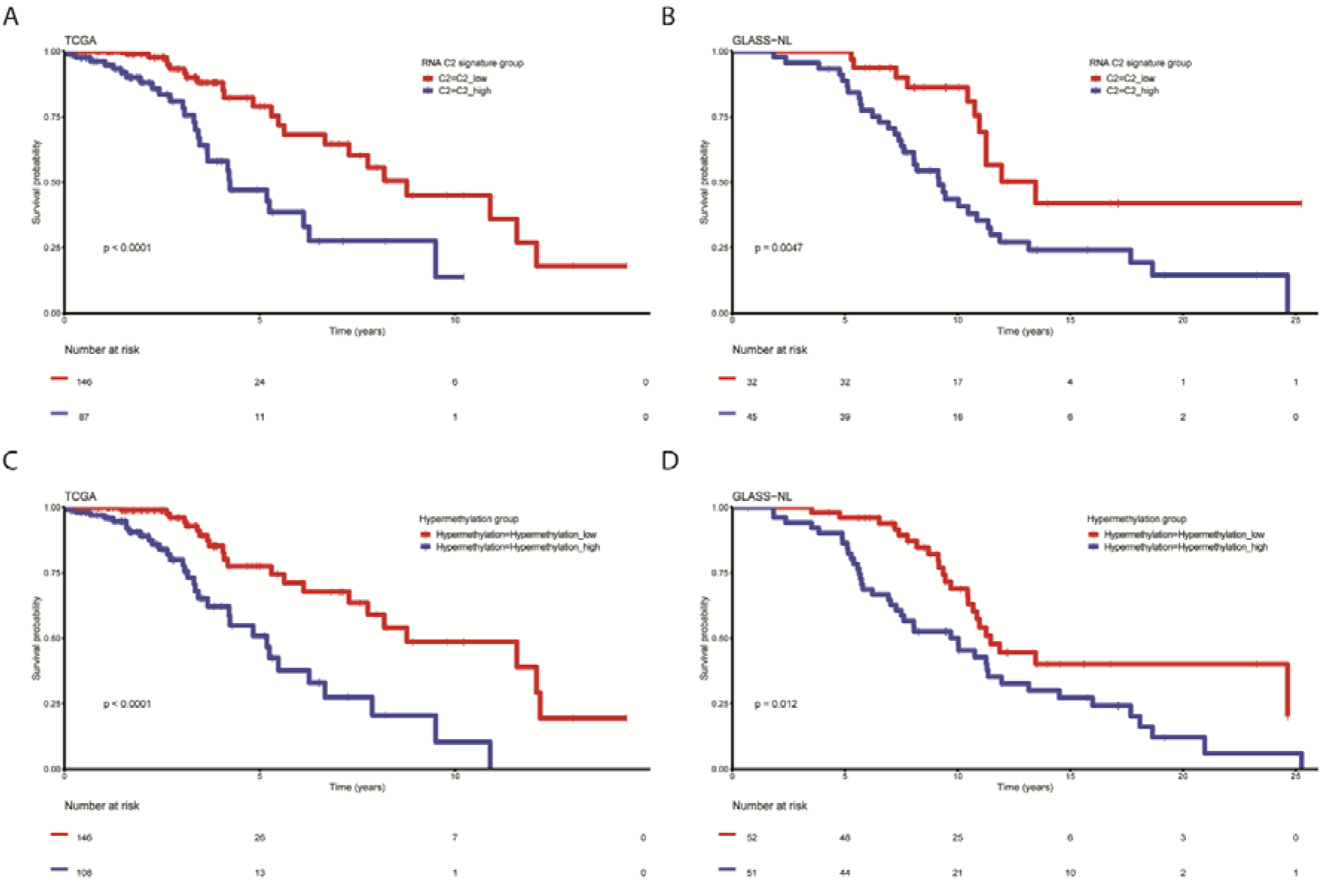
Kaplan-Meier overall survival curves of TCGA and GLASS-NL stratified by RNA C2 signature and hypermethylation risk groups.

## Notes

### Competing Interest Statement

P.M. Clement reports other support from EORTC during the conduct of the study; fees (to institution) from Bayer, Merck, Leo Pharma, Rakuten Medical, Takeda, and Bristol Myers Squibb (BMS); fees and nonfinancial support from MSD; and grants from AstraZeneca outside the submitted work; also fees for occasional advice to government agencies such as FAGG/EMA, as well as being a member of CTG (substitute) in Belgium. M.J. van den Bent reports consulting for Boehringer Ingelheim, F. Hoffman-La Roche, Fore Biotherapeutics, Genenta, Incyte Corporation, AnHeart therapeutics, Mundipharma, SymBio Pharma and Servier Affaires Medicales.

