## Supplementary Methods for "Epigenetic landscape reorganization and reactivation of embryonic development genes are associated with malignancy in IDH-mutant astrocytoma"

**DNA methylation**

In total 751 patients were enrolled in the CATNON trial (1,2). DNA was isolated from FFPE tissue as described earlier (3). After using the Infinium FFPE DNA Restoration Kit, DNA methylation profiling was performed with the Infinium MethlationEPIC BeadChip according to the manufacturer’s protocol. Patients were excluded for further analyses in case of insufficient material (n=78) or low-quality DNA methylation data (n=19). Low quality DNA methylation data was defined as >5% uninformative probes per sample at a detection P value < 0.01. 10 patients were classified as oligodendroglioma based on presence of 1p/19q codeletion and for 10 patients the *IDH1/2* status was unknown. 204 patients were excluded since tumors did not exhibit the *IDH1/2* mutation. The remaining 430 patients were included for this study.

Preprocessing of the raw idat files was performed using the minfi package (4). Background noise correction and dye-bias normalization were performed using the preprocessNoob function. Poor performing probes with a detection P value >.01 were omitted (CATNON: n=67,849, GLASS-NL: n=91,130, TCGA: n=38,435). Also, probes which were previously described to underperform were excluded (5) (Infinium EPIC array: n=105,454, Infinium 450K array: n=92,783).

**RNA-sequencing**
For the CATNON dataset, Illumina sequencing reads were obtained as compressed fastq files (n=183 samples). Unique Molecular Identifiers (UMI) were extracted to the read header using UMI-tools (v.1.2.2). Using fastp (v.0.23.2), low complexity reads were filtered, polyG reads longer than 35 bases were trimmed and unwanted polyX (i.e. PolyA) tailing was removed. After quality control with both fastp and fastQC (v.0.23.2), reads were mapped to the human reference genome (hg19/GRCh37) using the STAR aligner (v.2.7.9a). The minimum overhang for annotated and unannotated junctions were respectively set to 1 and 10 bases. Expected intron lengths were set between 20 and 1000000 bases. Maximum genomic distance between mates was set to 1000000. The minimum chimeric segment length and overhang for a chimeric junction were set to 12. Alignments that contained non-canonical unannotated junctions were filtered. Resulting bam files were sorted and indexed using SAMtools (v.1.9). UMI were deduplicated using umi tools whereby chimeric read pairs and unpaired reads were discarded. The template length was ignored for paired-end deduplication. The number of reads mapped to each chromosome was determined using SAMtools IdxStats. FeatureCounts (v.2.0.3) was used to count the number of reads for each corresponding gene using gencode v34 as gene annotation. The mean number of assigned reads was ~8.5 million.. Samples with FeatureCounts assigned read counts below 500,000 (n=28) were excluded. In case of duplicates (n=2) the sample with the highest number of assigned reads was selected. For samples with assigned read counts between 500,000 and 750,000 we excluded samples with a high percentage of reads mapping to alternate loci (>50%, n=2) and outliers in an unsupervised PCA (n=1). Two PCA outliers with more than 750,000 reads were excluded due to a bad score in multiple other QC metrics (alternate loci mapping, GC content and UMI count). 148 samples passed our quality control pipeline, from which we excluded re-resections (n=8) and samples without matching methylation data (n=2). Sequencing protocols and processing of read counts for the GLASS-NL dataset are described in an earlier study (6).

**Tumor purity estimation**

We evaluated two tumor purity estimation methods by their correlation with each other and with RNA expression levels of genes associated with neurons. One method was based on DNA Whole Exome Sequencing (WES) data to subsequently estimate tumor purity by the Variant Allele Frequency (VAF) of the IDH1/2 mutation. We hereby assumed no loss of heterozygosity and that the IDH1/2 mutation was a clonal mutation. The R-package InfiniumPurify estimates tumor purity by using informative differentially methylated CpG sites (iDMC) between tumor samples and a reference set of normal brain samples (7). The RNA expression levels per cell type were calculated by selecting the top 100 marker genes provided by McKenzie et al (8). This information was summarized to a single component per cell type, by using principal component 1 (PC1) from the output of Principal Component Analysis (PCA) on the marker genes per cell type.

**RNA-sequencing analysis of purified human CNS cell types**

We acquired transcriptome profiles (n=23,223 genes) of purified human CNS cell types from the original publication (9). In short, fetal astrocytes (n=6) were isolated from brain tissue samples obtained during elective pregnancy terminations, while mature astrocytes (n=12) were derived from juvenile and adult brain tissue samples (8y-63y) acquired during neurological surgeries. Oligodendrocytes (n=5) and tumor-associated macrophages/microglia (n=3) were isolated from adult brain tissue samples. We compared the mean Fragments Per Kilobase Million (FPKM) values for our downregulated (C0) and upregulated (C1-C3) transcriptional clusters between cell types.
